# Costs and benefits of maternal nest choice: tradeoffs between brood survival and thermal stress for small carpenter bees

**DOI:** 10.1101/2022.11.30.518597

**Authors:** JL deHaan, J Maretzki, A Skandalis, GJ Tattersall, MH Richards

**Author notes:** Corresponding author: MH Richards. Vineland Research and Innovation Centre, Biological Crop Protection, 4890 Victoria Avenue North, Vineland, Ontario, Canada (JL deHaan). Open Research Statement: Data are not yet provided *[Data will be archived at www.brocku.ca if the manuscript is accepted]*. Upon acceptance data will be archived in *[Brock University Respository, www.brocku.ca].

## Abstract

Nest site selection is a crucial decision for bees because where mothers construct their nests influences the developmental environment of their offspring. Small carpenter bees (*Ceratina calcarata*) nest in sun or shade, suggesting that maternal decisions about nest sites are influenced by thermal conditions that influence juvenile growth and survival. We investigated the costs and benefits to mothers and their offspring of warmer or cooler nest sites using a field experiment in which mothers and newly founded nests were placed in sunny or shady habitats. Maternal costs and benefits in sunny and shady treatments were quantified by comparing brood provisioning behaviour, nest size, number of brood cells, and offspring survival rates. Juvenile costs and benefits were quantified as body size, high temperature tolerance (CT_max_), metabolic rate, and pupal duration. The major maternal benefit of nesting in sun was significantly lower rates of total nest failure (caused by predation, parasitism or abandonment), which led to sun mothers producing 3.2 brood on average, while shade mothers produced only 2.9. However, sun nesting entailed costs to brood, which were significantly smaller, less likely to survive to adulthood and had significantly elevated CT_max_. This suggests that juvenile bees in sun nests bees experienced thermal stress during development, causing them to shunt resources from growth to thermoprotection, at the cost of smaller size and higher mortality. Pupae raised in a thermal-gradient “BeeCR” machine developed significantly faster at warmer average temperatures, which may be an additional benefit of sun nesting. Overall, our results highlight a tradeoff between maternal benefits and offspring costs when mothers choose nest sites, in which maternal fitness is enhanced by nesting in sun, despite significant physiological costs to offspring, due to the necessity for thermoprotective responses.

**Thinking through pandemic research:** The first lockdowns of the COVID-19 pandemic began as we prepared to enter the second field season of this study in 2020. Student research halted overnight. Lab access and travel were restricted. With limited access to field sites and no access to lab equipment, we brainstormed alternative approaches that would repeat, if not replicate, our main experiments of 2019 and fulfill degree requirements for JL de Haan’s MSc in a satisfying way. Our 2019 results had provided convincing evidence developmental temperature has long-term impacts on *C. calcarata* physiology, so we thought about which physiological measurements would be feasible outside the lab. Authors MH Richards and GJ Tattersall suggested collecting more measurements of CT_max_: the Peltier plate device required running water, but a portable water pump and a bucket allowed the apparatus to be set up anywhere. No calibration of instruments was required, and the only maintenance was to change the water in the bucket after a few hours of use. Thus, a student’s home basement became a laboratory. To investigate how temperatures affect developmental rate, we needed to raise bees in controlled environments, but incubators were not available. Author A Skandalis suggested repurposing a gradient PCR unit as a portable insect incubator (“The BeeCR”). The idea was tested successfully at home in 20202, so a larger study was done by J Maretzki in 2021 when undergraduate lab access was permitted again. Two outcomes of our pandemic pivot produced long-term benefits for our research. The BeeCR is a flexible, inexpensive, easy-to-use incubator perfectly suited for raising small insects at multiple simultaneous sets of variable temperatures. And the ease with which “field” sites could be established in our backyards demonstrates how amenable small carpenter bees are to field manipulations, suggesting this is a model species for addressing a variety of ecological and physiological questions.

## Introduction

For many animals, both vertebrates and invertebrates, nest site selection is a crucial prerequisite of raising offspring. Bees are central place foragers whose offspring must be raised in the shelter of a nest constructed in some sort of solid nesting substrate. Nests are crucial because juvenile bees are defenceless and mostly immobile, requiring considerable maternal care. In some bee groups, most notably the carpenter, orchid and sweat bees, mothers as well as offspring, spend much of their lives in their nests with their developing offspring, guarding them against predators and parasites, feeding young adults after they eclose (Hogendoorn et al. 2001; Quiñones and Wcislo 2015). In many bee species, offspring remain in their natal nests to overwinter and then reuse their nest to raise their own offspring – carpenter bees are known to reuse nests for many generations (Gerling et al. 1989; Cameron 2004; Schwarz et al. 2007). Thus, nest site selection has far-reaching fitness consequences for mothers and offspring, sometimes over multiple generations. Nest site selection in bees is influenced by both abiotic and biotic factors, including temperature, availability of nesting substrate, detectability of nests by predators and parasites, and proximity to food resources for both parents and offspring. Inevitably, choosing a nest site requires parents to evaluate multiple criteria, weighing the costs and benefits both for themselves as parents and for the welfare and survival of their offspring.

The best studied abiotic factor influencing nest site selection in bees is temperature (Potts and Willmer 1997; Potts et al. 2005; Hranitz et al. 2009; Antoine and Forrest 2021). Nests in sunny locations heat up earlier in the day and reach higher temperatures than do nests in shade. Since bees are ectotherms whose metabolism is closely linked to ambient temperature, the thermal microenvironment inside a nest influences everything from maternal behaviour to juvenile development. Nesting in a sunny location can be advantageous, because mothers may be able to warm up and forage earlier in the day, and larvae may develop faster. However, if nests in sunny sites heat excessively, then their occupants may experience heat stress, impairing development and survival (Chown et al. 2011; Corbet and Huang 2016; Wenzel et al. 2020). Nesting substrate not only influences the thermal environment of the nest, but also influences whether bees can respond to seasonal changes in temperature. For instance, ground-nesting bees can excavate brood cells at deeper soil depths when temperatures rise in summer, providing juveniles raised at different times of the year, with similar and optimal developmental temperatures (Antoine and Forrest 2021). In contrast, bees that nest above ground cannot usually adjust the size or position of their nests in response to seasonal changes in temperature from spring to summer. As a result, offspring produced in summer may experience conditions considerably warmer than were apparent to mothers selecting nest sites in spring.

Small carpenter bees excavate their nests in woody substrates like milled lumber or small twigs, and frequently choose sites with sunny exposures (Rehan and Richards 2010a; Vickruck et al. 2011). Field studies of small carpenter bees (genus *Ceratina*) demonstrate that females strongly prefer to nest in sunny sites, which provide internal nest temperatures warmer than ambient air temperatures (Vickruck and Richards 2012). Paradoxically, *Ceratina* mothers prefer a nesting substrate, raspberry (*Rubus)* plants, that grows most frequently in partial or complete shade. These intersecting site and substrate preferences suggest that *Ceratina* mothers must weigh the costs and benefits of nesting in preferred locations against the costs and benefits of nesting in preferred substrates. Vickruck and Richards (2012) carried out a field experiment to investigate the reproductive consequences of nesting in sun versus shade and in different types of substrate. *C. calcarata* mothers that nested in sun produced more and larger offspring, but so few bees chose shady sites that it was not possible to convincingly determine whether the apparent advantages of sun nesting were ascribable to sun *per se*, to differences in nesting substrate, or to some unmeasured biotic factor such as differential susceptibility to brood parasitism (Vickruck and Richards 2012).

One intriguing result in the study by Vickruck and Richards (2012) suggested that nesting in sun might have subtle physiological consequences for developing juveniles – this was the observation that when brought into the lab, juvenile bees from sun nests developed more slowly than those from shade nests. Since all bees experienced the same ambient temperatures in the lab, the difference implied that developmental conditions in sun and shade nests had induced long-lasting differences in offspring metabolism. To test this hypothesis, nests containing *C. calcarata* brood were moved into three thermal field treatments (full shade, semi-shade, and full sun), then tested to find out their metabolic rates at physiologically relevant temperatures (10, 25 and 40°C). The effect of rearing treatment became apparent at the highest temperatures, with bees from sun nests exhibiting the highest metabolic rates, those from shade nests exhibiting the lowest, and those from semi-shade nests showing intermediate metabolic rates. This indicated that bees that experienced higher daily temperatures inside their nests developed the ability to respond to higher temperatures by greater elevation of their metabolic rates. One adaptive consequence of nesting in sunny nests may therefore be the ability of juvenile bees to better cope with high temperatures, perhaps because their warmer microenvironments induce them to activate thermoprotective biochemical pathways, such as those involved in the production of heatshock proteins (Hofmann and Todgham 2010; Torson et al. 2017).

### Objectives

In this study we follow up on previous studies on the thermal microenvironment of Ceratina nests to examine tradeoffs inherent in maternal decisions to nest in warm, sunny sites versus cool, shady sites. We investigated consequences for both mothers and offspring, focussing on how nesting location affects maternal foraging activity and brood productivity, as well as physiological traits of brood, including body size, metabolic rate, and high temperature tolerance. We did this using a field experiment to manipulate the thermal microenvironments of nests by moving them into sunny or shady sites shortly after mothers established their nests, thus randomizing mothers among thermal environments to avoid possible maternal effects associated with nesting site decisions. To evaluate the consequences of sun versus shade nesting for mothers, we compared their foraging behaviour at nest entrances, as well as opening nests to compare brood productivity and survival. To evaluate the consequences of sun versus shade nesting for offspring, we allowed juveniles to develop to pupation under sunny and shady treatment conditions, then brought them into the lab to evaluate body size, metabolic rate, and temperature tolerance. Since previous experience showed that bees could not be raised successfully outside their nests under field conditions, we repurposed a gradient PCR machine as a variable temperature incubator for investigating pupal developmental rates over a wide range of temperatures. The onset of Covid-19 restrictions on travel in spring 2020, necessitated a pandemic pivot in which field work was moved to our backyards and a lab was set up in the first author’s basement.

## Methods

### Description of nest initiation and thermal treatment field sites

In 2019, nest initiation sites were set up in sunny, unmowed areas of the Brock University campus (St. Catharines, Ontario; 43.117°N, −79.256°W). We tied raspberry (*Rubus* sp.) canes cut to about 0.5 m in length to bamboo stakes and placed these in known areas with frequent nesting activity by *Ceratina calcarata* (Vickruck et al. 2011; Vickruck and Richards 2012; Lewis and Richards 2017a). Twigs were checked every two to three days prior to 0900 h. The founding of a nest by an adult female was confirmed visually by the presence of a female or sawdust in the nest entrance or by buzzing sounds emitted after gently inserting a blade of grass into the nest entrance (Lewis and Richards 2017). Nests with mothers were randomly assigned to a sun or shade treatment site and relocated to sun or shade sites on the same day. Before nests were moved, the entrance was temporarily blocked with tape to prevent the mother from escaping. There were two sun treatment sites in a meadow that received direct sun from 0800 h to 1800 h, and two shade sites inside the woodland margin on the edge of the same meadow that never received direct sun but did receive dappled sunlight in early morning and late evening.

In 2020, we used similar field methods but set up the nest initiation and treatment sites in different locations with different source populations of bees. Nest initiation sites were set up in sunny portions of private properties in the municipality of Lincoln (43.170°N, −79.481°W) and at the Glenridge Quarry Naturalization Site (GQNS; 43.121°N, −79.237°W). After nest initiation, nests were moved to treatment sites (one sun and one shade) in Lincoln. The sun site received full and direct sun from 0800 h to 1900 h, while the shade site received early morning sunlight prior to 0800 h and was shaded the rest of the day.

In 2021, nest initiation sites were set up at the GQNS and at a private property in Welland (43.002°N, −79.207°W) to provide juvenile bees for a lab study relating developmental temperatures and durations. There were no treatment sites.

### Temperature measurements in sun and shade treatments

We measured maximum daily nest temperatures using artificially constructed nest tunnels (“dummy nests”). These were created by drilling a hole 10 cm in length and 3.2 mm in diameter in the exposed pith of four raspberry canes. One dummy nest was placed at each of the treatment sites in 2019, and two were placed at each site in 2020. Daily maximum temperatures were recorded between 1300 h and 1500 h from 28 May to 20 August 2019 and from 3 June to 12 August 2020 using an Omega temperature unit (Model HH509R) by placing the wire thermocouple probe (T-type; 0.1°C precision) inside a dummy nest tunnel. The wire probe was held outside the nest at the height of the entrance to measure ambient air temperature. On rainy days it was assumed that nest temperature was equal to air temperature in both treatments, and air temperature was recorded from a covered patio near the treatment sites.

### Maternal foraging behaviour

Maternal foraging behaviour was quantified by observing adult females as they departed from and returned to nest entrances during the 2019 brood-provisioning period. Up to 10 randomly selected nests were observed throughout the course of a day (0800 h to 1800 h) on 16 days between 26 June and 26 July 2019. Observations were facilitated by enclosing nest entrances with inverted plastic cups and lids that prevented bees from exiting or entering their nests until the cup was removed; cup removals delayed bees by only a few seconds and bees rapidly became accustomed to the procedure (Lewis and Richards 2017). We distinguished between pollen flights (mother arrived at the nest with pollen visible on her hind legs) and nectar flights (mother arrived without pollen). We measured foraging activity in terms of the number of foraging trips (arrivals) per female per day, foraging trip duration (average time elapsed from a departure to the next arrival), and total daily foraging time (the summation of all flight durations per female per day) (Richards 2004; Lewis and Richards 2017).

### Collections of nests and their occupants

Based on observed nest initiation dates and observations of forager activity, we aimed to collect nests at the full brood stage, when mothers had completed brood provisioning and egg laying (Rehan and Richards 2010a). Nest entrances were covered with tape early in the morning, to ensure the mother was inside and brought to the lab (or house) to be opened. Nests were opened by splitting the twig lengthwise, and all nest occupants were counted as a measure of maternal reproductive success. Nests with live bee occupants were designated as active brood (youngest offspring was at the provision mass or provision mass with egg stages) or full brood (youngest offspring had reached the larval or pupal stage) (Rehan and Richards 2010a, 2013). Adults, pupae and larvae with uneaten provision masses were placed separately into 0.2 ml microcentrifuge tubes with a small hole in the lid. Juveniles were allowed to continue development until they were used for further experimental studies, reached adulthood or died. Burrow length was measured with a ruler.

### Traits of bees from sun and shade nests

#### Body size

Bees collected in 2019 and 2020 were weighed and measured. Wet mass of provision masses (including eggs), larvae, pupae and adult bees collected was measured immediately after removal from their nests. For eggs and larvae, wet mass included the mass of any pollen still attached. Dry mass was measured for all adults and for juveniles that died as pupae after drying the specimens in a 60°C oven for 48 hours. All masses were measured using an analytical balance (Mettler Toledo MX5, 0.00001g precision). In 2019, we measured both wet and dry mass, but due to lab closures in 2020, only dry mass was measured.

The head widths (HW) of adults or bees that died as pupae were measured across the widest portion of their face, including the compound eyes (Vickruck and Richards 2012). The costal vein length (CVL) of adults was measured starting from where the costal vein and radial vein diverge to the proximal edge of the stigma. Measurements were made to the nearest 33μm on an AmScope dissecting microscope at 30X using an ocular micrometer.

#### Metabolic rate, VCO_2_

We used a push flow-through respirometry apparatus to measure the CO_2_ output of bees at 25°C and 40°C. Only bees collected in 2019 were evaluated, as the apparatus was inaccessible during 2020 pandemic shutdowns. The intake air was first drawn through a container of expended Amsorb Plus (Armstrong Medical) by a PP-2 Dual Pump System V1.0 air pump on the low setting (Sable Systems International), to assist with removing incurrent CO_2_. We used a volumeter (2.0 L) to calibrate flow rate, and a Flo-Box controller and Mass-Trak flow regulator (Sierra Instruments) to regulate the flow rate to 50 ml/min. The incurrent air was then pushed through a Rota-meter flowmeter. Two of these flowmeters were used for direct flow rate comparisons near the beginning and end of the flow through system. These devices were compared twice daily to check for air leaks in the system. After the flowmeter, the incurrent air passed through a container of indicating Drierite (W.A. Hammond Drierite Co.), to remove water vapour, and another container of Amsorb Plus to fully remove any remaining CO_2_ from the incurrent air on its way to the insect. The CO_2_ produced by each subject was measured by a S151 CO_2_ Analyzer (Qubit Systems) in parts per million (ppm, set to 500 ppm range and calibrated with nitrogen gas). The S151 CO_2_ analyzer was zeroed each morning using the fine adjustment screw after running the apparatus for 20 minutes with CO_2_-free air.

A Pelt5 Temperature Controller was used to set the temperature of the chamber housing the bee (Sable Systems International). An infrared activity meter (AD-2; Sable Systems International) was used to monitor the activity level of each subject during the trials. CO_2_ production, chamber temperature, and activity level were recorded by the data acquisition software AcqKnowledge (MP100 Data Acquisition Interface and Universal Interface Module UIM100C, Biopac Systems Inc).

Individuals were first tested at 25°C and then at 40°C with 48 h of removal from the nest, and usually on the same day they were removed from their nests. Trials lasted 30 minutes, with five minutes at the start and end of each trial to capture a baseline CO_2_ value within the chamber, and a 20-minute observational period in which a single bee was added to the chamber. The formula for calculating the rate of CO_2_ production is: *V□co*_2_*= Flow Rate× (FeCO*2 *− FiCO*2) in which FeCO_2_ is the fraction of excurrent carbon dioxide, FiCO_2_ is the fraction of incurrent carbon dioxide, and *Flow Rate* is the incurrent gas flow rate (ml/min). We converted CO_2_ (ppm) to FCO_2_ by dividing by 10^6^.

#### Critical thermal maximum (CT_max_)

The critical thermal maximum (CT_max_) is an animal’s maximum functional temperature, or the point at which it loses organized motor control, and is a measure of its thermal tolerance (Lutterschmidt and Hutchison 1997; DeVries et al. 2016; Oyen and Dillon 2018). We compared CT_max_ for bees raised in sun and shade in 2019 and 2020. Bees were tested one at a time by adding a bee to a Peltier plate at 20°C and increasing the temperature at a rate of 1°C per min. CT_max_ was recorded as the temperature of the Peltier plate at the point when the bee could no longer maintain coordinated movement, characterized by falling over onto its back and being unable to right itself (Oyen and Dillon 2018). The bee was then immediately removed from the apparatus and the plate was allowed to cool to 20°C before the next bee was added. Bees that died within 30 minutes of testing were excluded from analysis since they may have surpassed their CT_max_. Most bees were tested the same day that they were removed from their nests.

### Effect of developmental temperature on pupal duration, Q_10_ and Arrhenius plot

To more precisely analyse the effects of warm versus cool developmental temperatures on developmental rate, we repurposed an Eppendorf Mastercycler PCR machine as an incubator (a “BeeCR” machine). We programmed a thermal gradient with temperature changes chosen to represent a range of 24-hour temperature profiles mimicking the coolest to warmest July days over the years 2012 to 2019 (Supplementary Table S1). Juvenile bees were collected from sunny nest initiation sites in 2021 and were placed into the BeeCR in individual 0.2 ml microcentrifuge tubes with the lids pierced to allow air exchange. Bees in the coldest part of the gradient (position 2) experienced average daily temperatures of about 12.7°C in position 2, whereas bees in the hottest portion of the gradient experienced an average temperature of 32.7°C in position 11. Bees placed in the middle of the gradient (position 6) experienced a daily average temperature of 21.3°C. For comparison, normal average temperatures in the region are about 19.0°C in June, 21.9°C in July, and 20.8°C in August (Environment Canada, St. Catharines A station, 1981-2020, https://climate.weather.gc.ca/climate_normals/). We recorded the developmental stage of each bee every 1-2 days using the 18 stages defined in Rehan and Richards (2010b). Bees that died before eclosion were replaced with new specimens. Developmental time was calculated as the number of days elapsed from putting a white-eyed pupa into the BeeCR until it eclosed as an adult.

We took the inverse of development time as the development rate and examined the temperature sensitivity of development rate in two ways. First, we calculated the Q_10_ for the sequential increments in temperature, employing the mean overall temperature for each position in the BeeCR experiments (Supplementary Table S1). Q_10_ was calculated as:

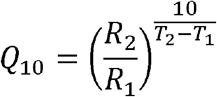

where R_2_ and R_1_ correspond to developmental rates at the corresponding temperatures, T_2_ and T_1_. Next, we used an Arrhenius plot to estimate the Arrhenius breakpoint temperature using the relationship between the log transformation of development rate and the inverse of absolute temperature (expressed in Kelvin^-1^) (Hansen et al. 2017). By examining whether there is more than one linear relationship describing temperature sensitivity, we can determine what critical temperature above/below which the rate of development suddenly shifts in its temperature sensitivity. We employed a two phase, one breakpoint regression model using the *segmented* package in R (Muggeo 2008).

### Additional data analysis

Data analysis and visualisation were carried out using R version 3.5.3 in R-Studio Version 1.2.5033. Regional weather data were downloaded from Environment Canada (St. Catharines A and Vineland Stations, https://climate.weather.gc.ca/climate_normals/). Ambient air temperatures on consecutive days are not independent, so to compare air and nest temperatures in sun and shade and between 2019 and 2020, we used Kolmogorov-Smirnov tests, which test whether two sets of numbers are drawn from the same distribution. For measures of nest size and brood productivity, we used two-way ANOVA models that simultaneously examined the effects of Treatment (sun vs. shade) and Year (2019 vs. 2020) and were initially of the form Variable ∼ Treatment * Year. Where interaction terms were not significant, we dropped the interaction. When multiple bees from the same nest were included in an analysis, we accounted for similarities among nestmates (which were often siblings) by using generalized linear mixed models with nest number as a random variable. For analyses of brood body size, we used generalized linear models in which brood sex, nest ID, and brood cell position were treated as random variables, since these variables influence brood body size (Rehan and Richards 2010c; Lawson et al. 2016).

## Results

### Thermal differences between sun and shade nests

The flight season of 2020 was significantly warmer than that of 2019 (Table 1, Supplementary Figure S1). In both years, sunny sites were warmer than shade sites during the hottest part of the day (Supplementary Figure S2). As a result, nests in sun reached maximum temperatures that averaged 1.6 to 3.8°C higher than in shade nests (Table 1).

**Table 1.** Comparison of ambient and nest temperatures for *Ceratina* nests in sun and shade during the warmest part of the day (14.00 h) in 2019 and 2020.

| Measurement (°C) | Thermal treatment | 2019<br>mean ± s.d.<br>(min, max) | 2020<br>mean ± s.d.<br>(min, max) | 2019 vs. 2020 |
| --- | --- | --- | --- | --- |
| Air temperature | Sun | 25.3 ± 4.0 °C<br>(13.7, 31.8) | 29.0 ± 3.9 °C<br>(20.2, 36.5) | **D=0.368, p~0.0002** |
| | Shade | 23.7 ± 3.7 °C<br>(12.8, 29.8) | 26.8 ± 4.2 °C<br>(17.4, 34.8) | **D=0.357, p~0.0003** |
|  | Sun vs. shade<br>difference | **1.6 °C**<br>**D=0.232, p~0.025** | **2.2 °C**<br>**D=0.254, p~0.044** |  |
| Internal nest<br>temperature | Sun | 27.0 ± 4.7 °C<br>(14.0, 34.0) | 30.8 ± 3.6 °C<br>(21.6, 37.3) | **D=0.362, p~0.0002** |
| | Shade | 24.2 ± 3.7 °C<br>(13.7, 30.2) | 27.0 ± 4.2 °C<br>(18.0, 34.8) | **D=0.326, p~0.0014** |
|  | Sun vs. shade<br>difference | **2.8 °C**<br>**D=0.366, p~3.42e-05** | **3.8 °C**<br>**D=0.458, p~8.61e-06** |  |
| Temperature<br>differential,<br>nest – air | Sun | 1.8 ± 1.2<br>(0, 4.1) | 1.8 ± 1.1<br>(0, 3.9) | D=0.137, p~0.538 |
| | Shade | 0.4 ± 0.4<br>(-0.3, 1.5) | 0.2 ± 0.2<br>(-0.3, 0.8) | **D=0.372, p~0.0002** |
|  | Sun vs. shade<br>difference | **1.4 °C**<br>**D=0.610, p~1.15e-13** | **1.6 °C**<br>**D=0.847, p~2.2e-16** |  |

Notes: Air and nest temperatures were measured during the hottest part of the day (~14.00h). Kolmogorov tests (D) were used to compare paired temperature series by date (air versus nest temperatures or 2019 versus 2020). The higher temperature differentials for nests in sun indicate that they warmed up more than nests in shade. Nests were also warmer in 2020 than in 2019.

### Influence of sun and shade nesting on maternal behaviour and reproduction

We examined the pollen and nectar-collecting behaviour of mothers following nest relocation to sun or shade treatments in 2019, based on three parameters, the number of foraging trips per day, the duration of foraging trips, and total flight time per day. Sun and shade mothers did similar amounts of pollen foraging (Figure 1). However, they differed in terms of nectar foraging, with sun mothers doing more nectar trips and spending considerably more time foraging for nectar on a daily basis.

**Figure 1.**
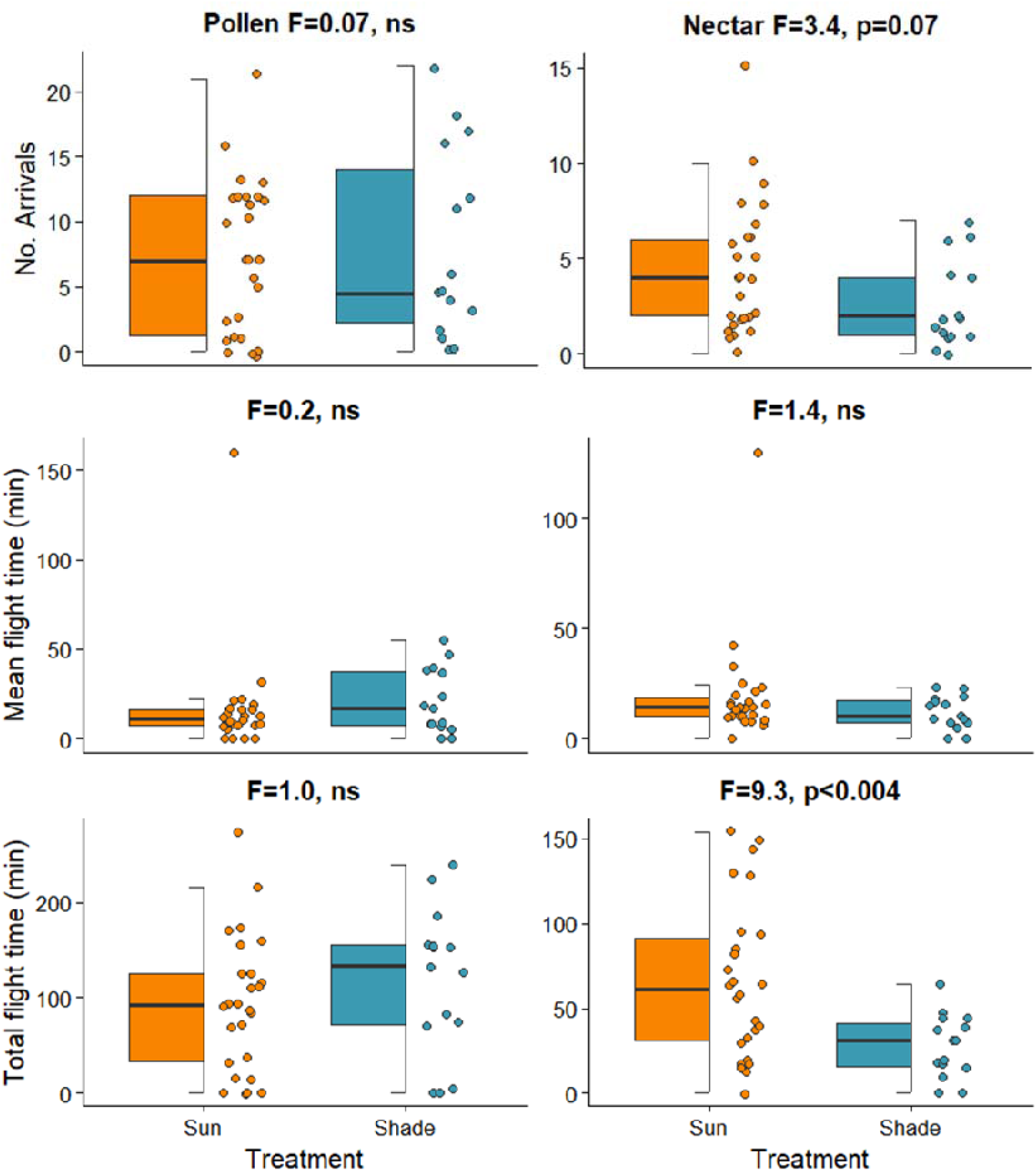
Comparison of pollen and nectar foraging metrics between sun and shade mothers in 2019. Results of ANOVA tests (with df=1,39) comparing daily foraging activities of females in sun and shade are given in each panel. Points represent individual mothers.

The contents of 194 nests were examined in 2019 and 2020. Nests in sun were significantly more likely to contain brood (categorized as full brood or active brood), whereas nests in shade were significantly more likely to have experienced total nest failure (no live brood) (Table 2). Overall, similar proportions of nests failed in 2019 and 2020, but it was mostly impossible to discern the reasons for nest failure.

**Table 2.** Status of all *Ceratina calcarata* broods when nests were collected in 2019 and 2020.

| Status of nest when collected | 2019 |  |  | 2020 |  |  |
| --- | --- | --- | --- | --- | --- | --- |
|  | Sun | Shade | Total | Sun | Shade | Total |
| Full brood | 25 | 16 | 41 | 15 | 7 | 22 |
| Active brood | 9 | 4 | 13 | 7 | 5 | 12 |
| Empty | 22 | 34 | 56 | 15 | 30 | 45 |
| Lost or predated | 1 | 1 | 2 | 1 | 2 | 3 |
| Total no. nests | 57 | 55 | 112 | 38 | 44 | 82 |
| Nests with brood | 34 (60%) | 20 (37%) | 54 | 22 (58%) | 12 (27%) | 34 |
| Nests without brood | 23 (40%) | 35 (60%) | 58 | 16 (42%) | 32 (63%) | 48 |
| Chi-test | **X<sup>2</sup> = 5.19, df = 1, p = 0.023** |  |  | **X<sup>2</sup> = 6.67, df = 1, p = 0.010** |  |  |

Notes: Nest fate is the status of the nest at the time of collection. Empty nests contained neither brood nor an adult female (a mother) and could have been predated or abandoned. Significantly more sun than shade nests contained brood at the time of collection, indicating that total nest failure was more frequent in shade. The overall proportions of nests with and without brood were similar in 2019 and 2020 ( $X^2=0.87$ , $df = 1$ , ns).

To compare maternal brood productivity in sun and shade, we focused on full brood nests in which mothers had ceased brood provisioning (Table 3). Within years, sun and shade nests had tunnels of similar length and contained similar numbers of brood cells. However, there were significantly more live brood in shade nests. Nests collected in 2020 were longer and contained more brood cells than those from 2019 but contained similar numbers of live brood. The sex ratio of brood that survived to the pupal stages was similar in sun and shade nests.

**Table 3.** Nest and brood conditions for full brood nests in sun versus shade.

| Variable | 2019<br>(cool) |  | 2020<br>(warm) |  | Treatment effect<br>(sun vs. shade) | Year effect |
| --- | --- | --- | --- | --- | --- | --- |
|  | Sun | Shade | Sun | Shade |  |  |
| N (nests) | 25 | 16 | 15 | 7 |  |  |
| Tunnel length (cm) | 13.6 ± 3.3 | 15.2 ± 3.5 | 15.9 ± 3.4 | 16.2 ± 3.3 | **F=1.34, n.s.** | **F=4.57, p=0.037** |
| No. brood cells (C) | 5.5 ± 2.7 | 7.7 ± 3.1 | 9.5 ± 5.1 | 10.0 ± 1.9 | **F=1.96, n.s.** | **F=12.81, p=0.001** |
| No. live brood (D) | 4.9 ± 2.6 | 6.6 ± 3.2 | 5.9 ± 3.4 | 8.4 ± 2.9 | **F=5.89, p<0.018** | **F=2.73, n.s.** |
| Sex ratio<br>(% male) | 67% | 48% | 56% | 68% | 2019: $X^2 = 0.10$,<br>n.s.<br>2020: $X^2 = 1.56$,<br>n.s. | |

Notes: F values represent partial F statistics with 1 degree of freedom, based on 2-way ANOVA (model statement: variable ~ Treatment + Year). Chi-square tests represent 2x2 contingency tests on total numbers of male and female brood collected each year, with 1 degree of freedom.

### Influence of sun and shade nesting on juvenile bee size and metabolism

Body size measures of juvenile bees collected as larvae, pupae or imagoes (from both full and active broods) are compared in Figure 2. After accounting for offspring size variation due to sex, brood cell position and maternal size, bees from sun nests tended to be smaller than those from shade nests in terms of linear measurements (head width and costal vein length) but not in terms of mass (wet and dry mass; Figure 2, Table 4). Dry masses of offspring from 2020 nests were significantly larger than those from 2019 nests, but they did not differ significantly in terms of head width or costal vein length (Table 4).

**Table 4.** Treatment (sun versus shade) and year (2019 versus 2020) effects on four measures of brood body size, based on the partial effects from generalized linear models in which brood sex, cell position, and family ID were treated as random effects.

| Response variable | Predictor variables |  |
| --- | --- | --- |
|  | Treatment<br>(sun vs. shade) | Year<br>(2019 vs. 2020) |
| Head width (mm) | **Shade > Sun:**<br>**F=6.98, df=1,55.7, p=0.011** | Similar:<br>F=1.96, df=1,57.3,n.s. |
| Costal vein length (mm) | **Shade > Sun:**<br>**F=6.04, df=1,51.2, p=0.017** | Similar:<br>F=2.52, df=1,52.6, n.s. |
| Wet weight (g) | Similar:<br>F=1.54, df=1,33.7, n.s. | - |
| Dry weight (g) | Similar:<br>F=2.11, df=1,65.6, n.s. | **2020 > 2019:**<br>**F=13.03, df=1,66.0, p=0.0003** |

Notes: These analyses were based on a general linear model of the form: Size measure ~ Treatment x Year. No interaction effects were significant and so were dropped from the final model. For wet weight, the Year term was excluded from the model since measurements were not available for 2020.

**Figure 2.**
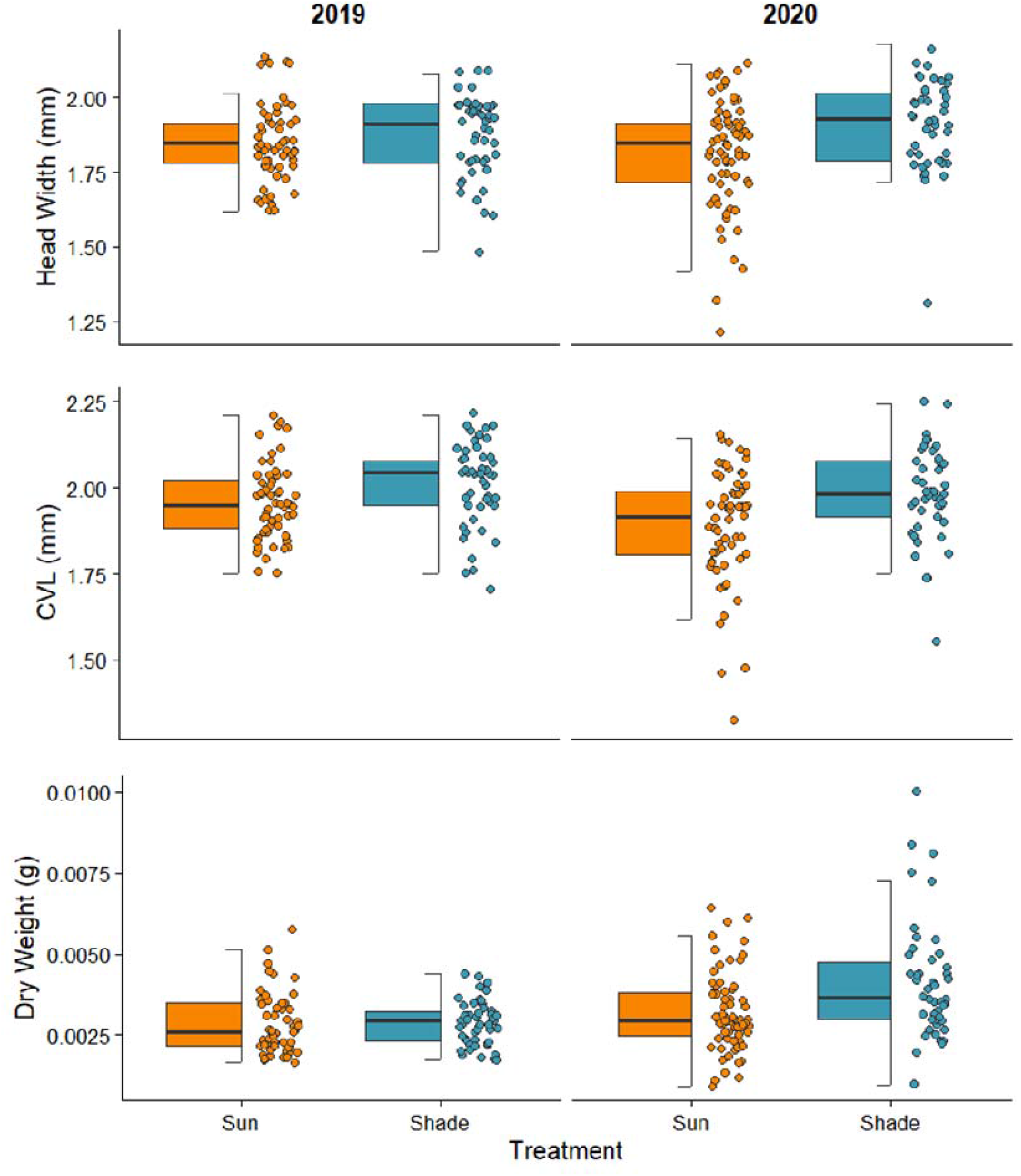
Body size measures for female brood collected as larvae, pupae or newly eclosed adults. Overall, brood were smaller in sunny than in shady nests. Results for male brood were the same. Note that wet weight of brood could not be measured in 2020, because of limited pandemic lab access, and so is not shown. Statistical comparison given in Table 4.

Variation in juvenile metabolic rate, as measured by VCO_2_, was significantly associated with bee mass, test temperature (25°C < 40°C), developmental stage (early pupa, late pupa, or newly eclosed adult), but not with field treatment (sun or shade) or sex (Figure 3; Table 5). CT_max_ was significantly higher for bees from sun nests and for bees collected in 2020 (Figure 4).

**Table 5.** Generalized linear model examining variation in metabolic rates of *Ceratina calcarata* raised in sun and shade nests (conditional R^2^ = 0.346).

| Predictor variable | NumDF | DenDF | F | p |
| --- | --- | --- | --- | --- |
| Mass | 1 | 182 | 14.59 | **<0.001** |
| Test Temperature (25°C and 40°C) | 1 | 182 | 67.24 | **<0.001** |
| Treatment (Sun vs. Shade) | 1 | 182 | 0.25 | 0.620 |
| Sex | 1 | 182 | 0.51 | 0.471 |
| Stage (Early pupae, late pupae, adults) | 2 | 182 | 10.50 | **<0.001** |
| Test Temperature * Treatment | 1 | 182 | 0.13 | 0.724 |

Notes: Since each individual was tested at both temperatures, individual bee ID was set as a random variable. Abbreviations: NumDF, numerator degrees of freedom; DenDF, denominator degrees of freedom.

**Table 6.** Fitness consequences for mothers and offspring of nesting in sunny and shady microenvironments, as well as comparisons between two years with significantly different weather.

| **Fitness measure** | **Warm site (sun) vs.**<br>**Cool site (shade)** | **Warm year (2020)**<br>**vs. Cool year (2019)** |
| --- | --- | --- |
| Maternal foraging effort - pollen | = |  |
| Maternal foraging effort – nectar | Warm > Cool | n.a. |
| Incidence of total nest failure | Warm < Cool | = |
| Tunnel length in *fb* nests | = | Warm > Cool |
| Brood cells in *fb* nests | = | Warm > Cool |
| Live brood in *fb* nests | Warm < Cool | = |
| *Brood survival in fb nests* | *Warm < Cool* | *Warm < Cool* |
| Brood body size (HW, CVL) | = | = |
| Brood body size (mass) | Warm < Cool | Warm < Cool |
| High temperature tolerance (CTmax) | Warm > Cool | Warm > Cool |
| *Thermal stress* | *Warm > Cool* | *Warm > Cool* |
| Metabolic rate | = | n.a. |
| *Development time* | *Warm < Cool* | *Warm < Cool* |

Notes: the performance comparison between 2019 and 2020 is confounded by the fact that bees were sampled from different geographic populations each year, due to pandemic constraints on travel in 2020. Traits denoted in italics were inferred, rather than directly measured, as described in the text. Abbreviations: *fb*, full brood.

**Figure 3.**
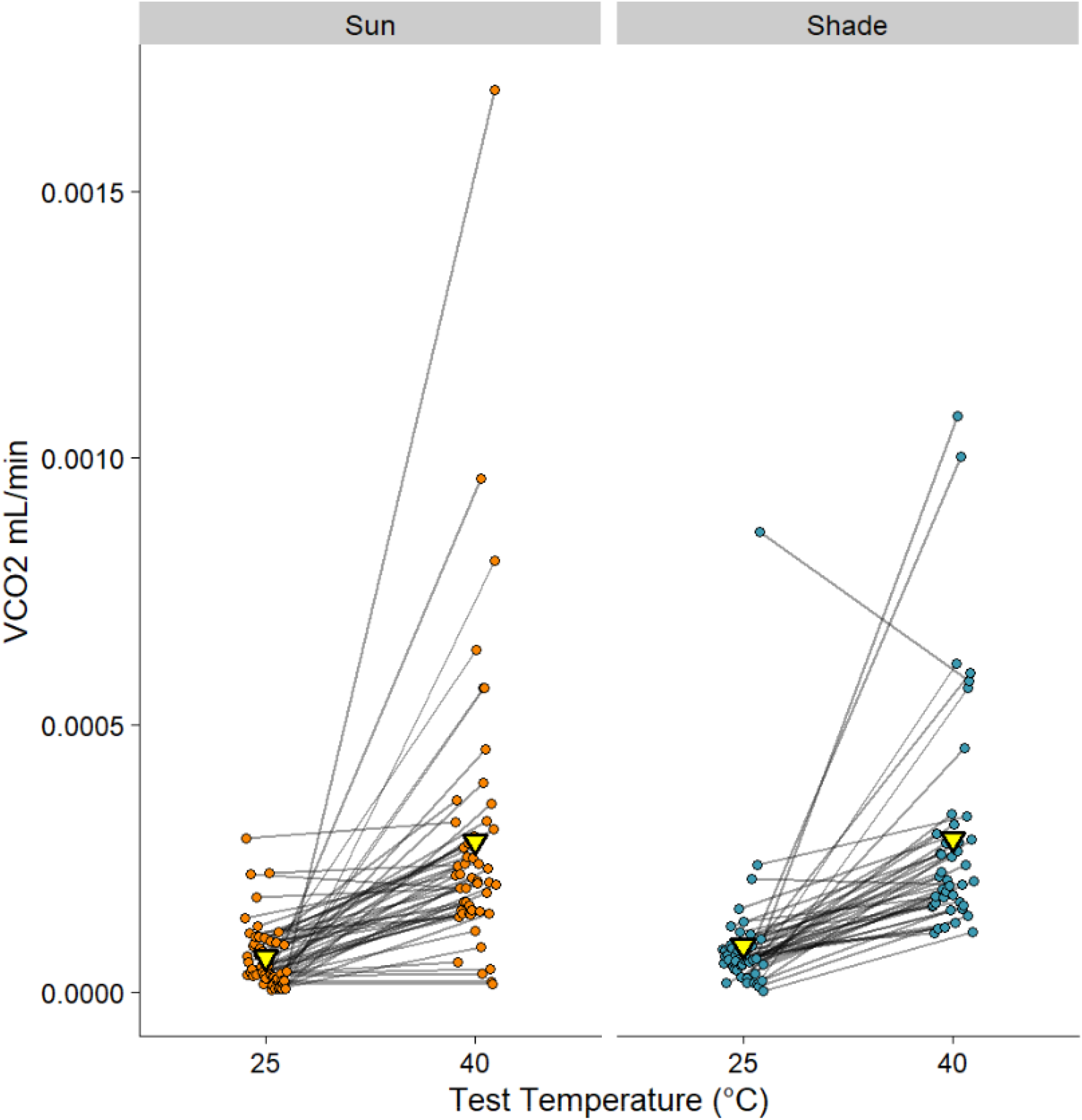
Bees from sun and shade nests had similar average metabolic rates. Each bee was tested at two temperatures and is represented by two points joined by a line. The mean VCO2 for each treatment at each test temperature are shown with yellow triangles. See Table 5 for statistical analysis.

**Figure 4.**
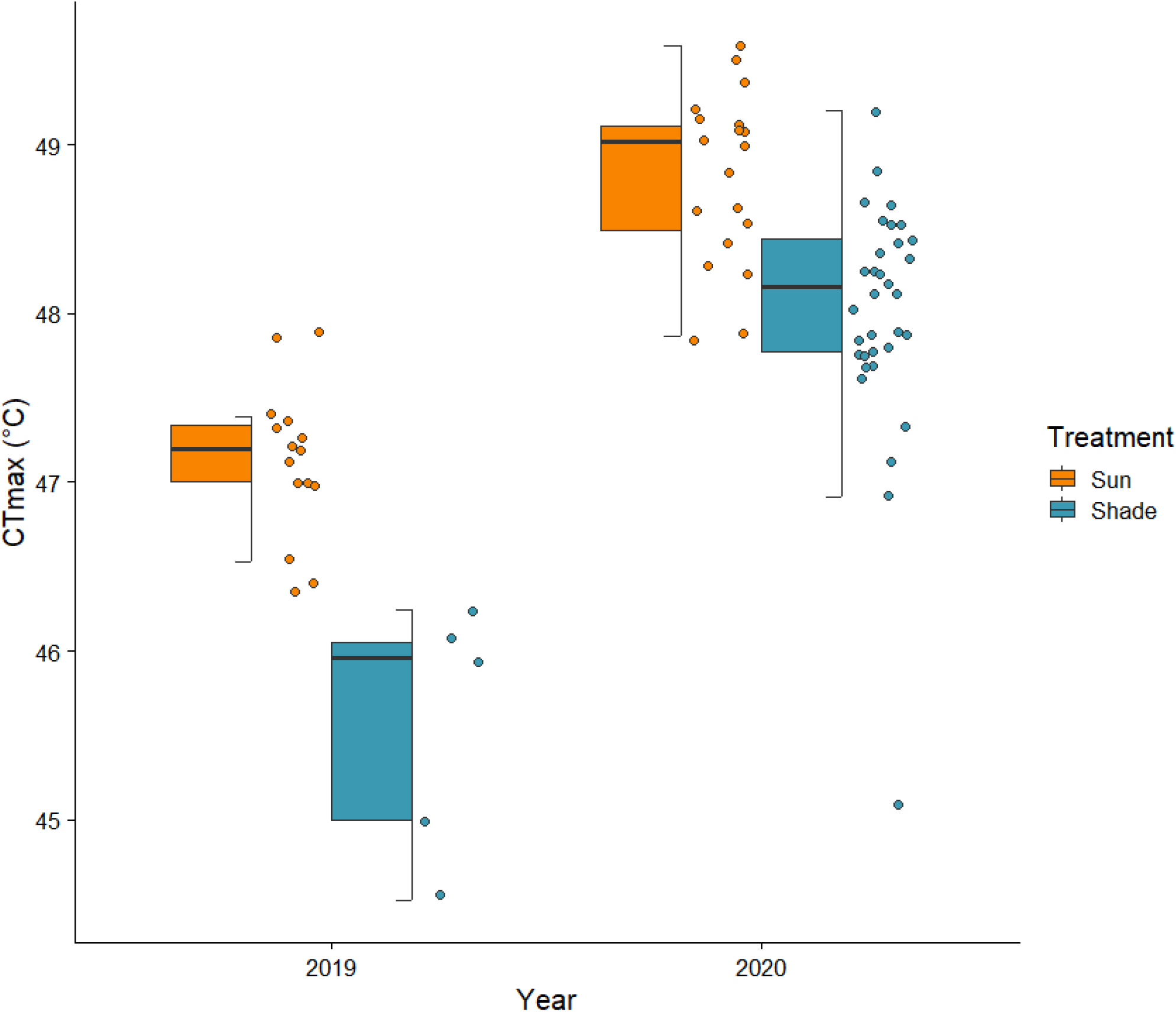
High temperature tolerance of newly eclosed adult *C. calcarata* from sun and shade nests in 2019 and 2020 as measured by CT_max_. Note that 2020 was a significantly warmer year than 2019.

### Influence of temperature on pupal developmental rate and thermal sensitivity, Q_10_

The developmental time of bee pupae from the white-eyed stage to adult eclosion was inversely correlated with the mean daily incubation temperature, with bees raised in the coldest temperatures developing about four times slower (∼60 days) than those at the warmest temperatures (∼15 days; Figure 5). An Arrhenius plot of developmental rate versus temperature (in degrees Kelvin^-1^) indicated a breakpoint temperature of about 19.6°C (95% CI: 18.4-20.8°C). A plot of Q_10_ versus temperature indicates extreme thermal sensitivity at low temperatures, with a decline in sensitivity towards Q_10_ values of about 2 or 3 between 21 and 32°C, falling below 1 for the highest temperature range studied (31.4 to 32.8°C).

**Figure 5.**
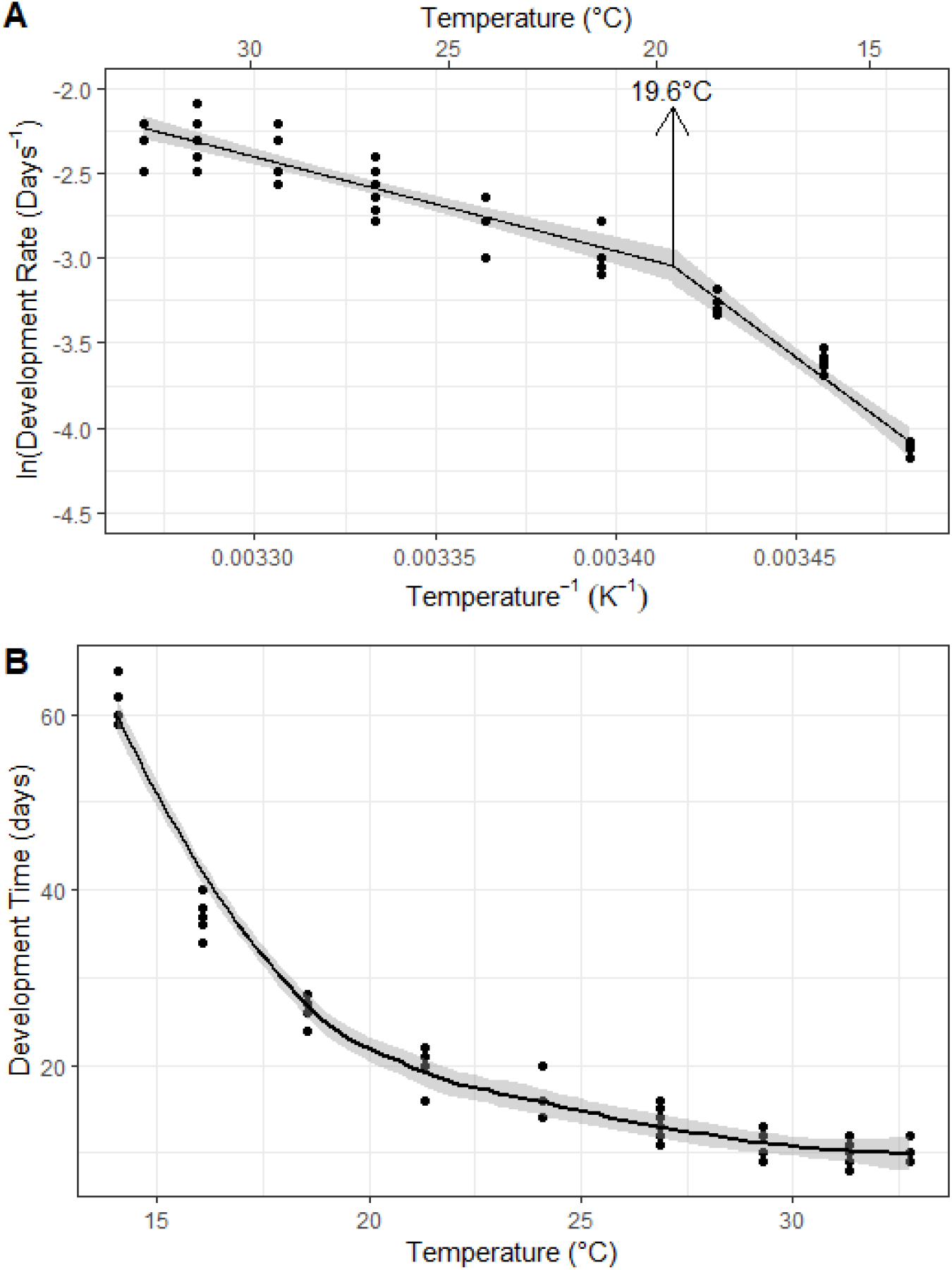
Relationship between average developmental temperature and pupal developmental rate from white-eyed to adult stages of *Ceratina calcarata* pupae raised in BeeCR thermal gradients, depicted as A) an Arrhenius plot of ln(Development Rate) vs. Temperature^-1^ showing a break-point temperature of 19.6 °C, and B) the Development Time with a smooth curve (± se in shading). Notes: A; The influence of average developmental temperature on pupal development rate. B; Developmental times for *Ceratina calcarata* pupae. Pupae were placed in the BeeCR at the white-eyed stage and removed as fully eclosed adults. Temperatures are the mean daily temperatures over a 24 h period, as presented in Table S1.

## Discussion

### Maternal reproductive outputs in sun and shade

Bees provision their offspring with two basic food resources, pollen and nectar. Our observations of carpenter bee mothers at nest entrances demonstrated that shade and sun mothers expended similar effort foraging for pollen, whereas sun mothers expended significantly more effort (measured by total daily flight time) foraging for nectar. Mothers likely feed themselves outside the nest, so pollen and nectar provisions brought back to the nest would be destined for larval consumption. The extra time spent on nectar trips by sun mothers would have allowed them to add proportionately more nectar to larval provision masses. In sun nests, both provision masses and larvae would be more susceptible to desiccation than in shade nests. Provision masses with more nectar would supply more liquid to larvae and might humidify the brood cells. In honeybees, the proportion of foragers returning with water and nectar increases as temperatures rise (Cooper et al. 1985).

There may be additional consequences of adding extra nectar to larval provisions in sun nests. Larval growth and performance may be influenced by the sugar content of the nectar added to their provision masses. Leaf-cutter bee larvae whose pollen masses are supplemented with extra sugar grow to larger sizes and sometimes develop more slowly than those without (Burkle and Irwin 2009). If carpenter bees followed this pattern, we would expect that brood from sun nests would be larger in size than brood from shade nests. In fact, we observed the opposite pattern, so extra nectar added to larval provisions in sun nests did not produce larger offspring. The smaller size of sun offspring is at odds with the observation that sun and shade mothers exhibited similar rates of pollen foraging, which should have produced offspring of similar body sizes (Danforth 1990; Bosch and Vicens 2002). That sun offspring ended up smaller suggests that during development, they diverted energetic resources away from processes promoting growth and so ended up smaller (Atkinson 1994; Richards et al. 2020).

A greater proportion of nests in shade were empty when collected, containing neither brood nor an adult female. This means that the sun mothers had a higher probability of successfully raising at least one offspring than did shade mothers. In the majority of empty nests, we could not determine what had happened to the occupants, but it is possible that being relocated from sun to shade led to higher rates of abandonment by shade mothers. Bee brood are helpless to defend themselves against predators and parasites, and nests left unguarded after a mother dies or abandons it are quickly raided by ants and other predators, which kill and remove all brood (Lewis and Richards 2017). In our experiment, initial nest founding by mothers was in sunny sites, after which we randomly assigned each nest to sun or shade treatment, so it is possible that mothers were more likely to abandon their nests if they were moved to shade. Another possibility is that nest failures were due to higher predation or parasitism, rather than maternal abandonment. Shade nests are more attractive and vulnerable to parasites, especially if constructed in a native plant substrate (Vickruck et al. 2011; Vickruck and Richards 2012). We noticed that some of the failed nests had short (<10cm) tunnels that contained eggs or developing larvae of tree crickets (*Oecanthus* sp.), a known pest of raspberry plants (OMAFRA 2009). Additionally, sun nests might have experienced lower rates of nest failure because parasites avoided them. This could happen if host nests heat to higher temperatures than can be tolerated by parasite larvae, a situation that seems to explain why red mason bees (*Osmia rufa)* reared at higher temperatures have fewer parasites than those reared at lower temperatures (Giejdasz and Fliszkiewicz 2016).

Among surviving nests, shade nests contained more live offspring and sun nests produced fewer. Since tunnel length, the number of brood cells in surviving nests, and maternal pollen foraging rates were all similar in sun and shade, fewer live offspring in sun nests implies higher mortality for sun brood during development. In other words, mothers nesting in sun less frequently experienced catastrophic nest failure, but apart from this source of mortality, fewer of their brood survived to adulthood. In contrast, mothers nesting in shade were more likely to lose their entire brood, but if they avoided this, then more of their brood survived.

We can estimate the average reproductive success of sun and shade mothers by applying the simplifying assumption that the rates of total nest failure and brood productivity in Tables 2 and 3 respectively, accurately represent population level rates. If so, then in sun nests, 41% of mothers produced 0 brood while 59% produced an average of 5.4 brood, producing an estimate of about 3.2 brood per mother. In shade nests, 62% of mothers produced 0 brood while 38% produced an average of 7.5 brood, producing an estimate of 2.9 brood per mother. These simple calculations suggest that for sun mothers, lower rates of total nest failure more than compensated for lower brood survival, such that sun mothers produced more brood than shade mothers.

This demographic benefit of sun nesting must be weighed against possible costs to offspring of developing at higher temperatures. One of these costs is smaller body size, as sun brood were smaller than those from shade nests, despite probably having consumed more sugar, which should have resulted in larger body sizes. In bees, smaller body sizes generally are associated with lower survival and reproductive success in both sexes (Shimamoto et al. 2006; Greenleaf et al. 2007; Skandalis et al. 2009; Rodríguez-Gironés and Bosch 2012; Grula et al. 2021). In *Ceratina* specifically, smaller females are less likely to survive overwintering hibernation (Rehan and Richards 2010a) and as mothers, they produce fewer and smaller offspring (Rehan and Richards 2010c). Smaller males may have lower mating success and perhaps lower survival (Alcock 1997). Long-term costs of smaller body size would suggest that the calculations above somewhat overestimate the advantages of nesting in sun. Nevertheless, sun nesting would appear to be the higher fitness strategy.

### Physiological responses of offspring to sun and shade microenvironments

The CT_max_ of adult offspring from sunny nests was about 0.8°C higher than that of shade bees, which demonstrates that their warmer microenvironments during development had long-term effects on how they respond to high temperatures. The observed CT_max_ of both shade and sun bees averaged about 48.3°C, which closely matches the reported CT_max_ =48°C of another *C. calcarata* population, as well as several tropical *Ceratina* species (Hamblin et al. 2017; da Silva et al. 2021). This value of CT_max_ is about 10°C higher than the maximum temperatures inside nests on warm days, so in our study, both shade and sun bees should have been inside their thermal safety margin (Sinclair et al. 2016). However, larvae and pupae might have different thermal performance curves than adults, with lower CT_max_ and perhaps lower thermal optima. Nevertheless, if bees were experiencing temperatures above their thermal optima during development, we would expect declining performance and the necessity to mount a response to thermal stress.

As expected, *C. calcarata* offspring exhibited higher metabolic rates (VCO_2_) at 40°C than they did at 25°C, but sun and shade bees did not differ in their responses to temperature. The similarity in response contrasts with a previous study, in which bees from sunny nests showed elevated metabolic rates at 40°C (Richards et al. 2020). Fieldwork for the previous study was carried out during the summer of 2016 which was one of the hottest, driest summers on record in southern Ontario, with temperatures slightly warmer than in 2020 (Supplementary Figure S2). This suggests that thermal differences between shade and sun treatments were large enough that year to induce a detectable change in metabolic responses of sun bees to high test temperatures, whereas in 2019, which was a moderate summer, conditions in sun nests were not extreme enough to induce the same response observed in 2016. Since 2020 was almost as warm as 2016, we suspect that we would have detected elevated metabolic rates of sun bees to high test temperatures in 2020, if pandemic shutdowns had not prevented those studies.

The developmental progress of juvenile bees could not be observed under field conditions, because outside of their nests, they were rapidly eaten or succumbed to weather-induced damage (J. de Haan, pers. obs). Instead, we used the BeeCR to raise *Ceratina* pupae in a gradient of daily temperature cycles that included temperatures such as those in sun and shade nests. Combining this information with local temperature records, we can estimate developmental rates in sun and shade, assuming that sun nests reached maximum temperatures about 4°C higher on average than shade nests (this study, Richards et al. 2020). In 2019, the mean air temperature at our research sites was 20.5°C, which predicts a pupal duration of 19.8 days in shade and 15.3 days in sun. In 2020, the temperature was about 22.6°C which predicts a pupal duration of about 17.3 days in shade and 13.4 days in sun. If these predictions are realistic, then pupae in sun nests develop about 4 days faster than those in shade nests. This represents a significant shortening of the window of opportunity for parasites and predators to consume developing bee brood and may be an explanation for the lower rates of total nest failure in sun.

Bees raised at different temperatures demonstrated enormous variation in pupal developmental time, from 8 to 65 days, far more variation than would ever occur in natural conditions. In the BeeCR, the warmest temperature profiles were about 10°C higher than the average profile, with a high of 37.3°C and a low of 29.4°C, reflecting temperatures on the warmest summer days recorded in Niagara between 2012 and 2019. The coolest temperature profile had a high of only 16.8°C, a low of 9.7°C, and a mean of 12.2°C, reflecting the coolest summer day recorded in Niagara during the same time period. What was unrealistic, was that pupae experienced these hot and cold extremes repeatedly, day after day. In Figure S3, the breakpoint in thermal sensitivity occurs around 21°C, which happens to be close to the mean developmental temperature under field conditions. The greater sensitivity of pupae to temperatures below this threshold is interesting since they would rarely experience such temperatures during development. The increased thermal sensitivity of development at these cooler temperatures must reflect the fact that developmental rate is optimised for much warmer temperatures. For temperatures above the breakpoint, we see clear, but fairly normal biochemical sensitivity of development rate (Q_10_ ∼2-3), eventually reaching a temperature where development actually slows down (∼32°C) reflective of the Q_10_ falling below 1. Increases in temperature above 32°C are likely to result in larvae failing to develop.

### The cost of thermoprotective responses

For juvenile bees in sun nests, three important consequences of development at the higher temperatures experienced in sun nests were smaller body size, higher mortality, and an upwards shift in CT_max_. Ectotherms, including bees, that are exposed to extremely high, but sublethal temperatures can mount a protective heat shock response if they do not warm up too fast (Hranitz and Barthell 2003; Elekonich 2009; Hranitz et al. 2009). The combination of higher CTmax, smaller body size, and higher brood mortality in brood from sun nests, suggests that these juveniles experienced sufficient thermal stress to necessitate a thermoprotective response, in which developmental resources were diverted from growth to survival. Heatshock responses to acute thermal stress have been observed in multiple insect groups (Krebs and Feder 1997; Hranitz and Barthell 2003; Elekonich 2009; Torson et al. 2017; Zhu et al. 2017). Heat shock proteins (HSPs) protect other cellular proteins from becoming damaged and promote recovery after the heat stress (Hofmann and Todgham 2010). Some HSPs are upregulated after an extreme thermal event or after chronic exposure to stressful temperatures, to protect the organism against other such events (Kim et al. 2017). Overexpression of the heat-shock response results in a decrease in growth and development in *Drosophila melanogaster* (Krebs and Feder 1997) and is associated with higher mortality in heat-exposed juvenile leafcutter bees (Hranitz et al. 2009). From the point of view of both mother bees and their offspring, thermoprotection is a cost of nesting in sunny sites, but evidently it is a cost that for mothers, at least, is more than balanced by the advantages of higher *per capita* reproduction (Hranitz and Barthell 2003; Hranitz et al. 2009).

### The functional significance of CT_max_

CT_max_ is a convenient and apparently simple measure of maximum heat tolerance (Jorgensen et al. 2021). A simplistic interpretation of CT_max_ would suggest that thermal differences between sun and shade nests were too small to matter, since the average CT_max_ of 48°C implies that bees in both sun and shade would be well inside their thermal safety margin (Sinclair et al. 2016). Yet the upward shift in CT_max_ due to larval experience in sun nests suggests that CT_max_ provides a useful measure of the sun bees’ response to thermal stress. Recent studies suggest that the most commonly measured acute CTmax is systematically higher than a CTmax measured over longer time frames. In other words, the effects of exposures to warm stressors accumulate over time, and thus chronic exposure to less severe extreme temperatures could still be lethal for many ectotherms (Jorgensen et al. 2021). Evidence is accumulating that intra-specific variation in CT_max_ is considerably greater than interspecific variation, implying that intraspecific variation is an important phenotypic indicator of physiological responses to thermal stress (Sinclair et al. 2016). In fact, evidence is accumulating that acclimation at higher temperatures results in higher heat tolerance (Jorgensen et al. 2021). Nevertheless, higher heat tolerance likely has a cost. In our study, the higher CT_max_ of sun bees was linked with higher brood mortality and smaller body sizes, suggesting that improved thermal tolerance in surviving brood was achieved by diverting resources from growth towards thermoprotective functions at the cost of reduced growth efficiency and higher mortality (Richards et al. 2020). Although there seems to be little variation in CT_max_ at the species level in *Ceratina* (da Silva et al. 2021), we would argue that the current study demonstrates that intraspecific variation in CT_max_ has functional significance.

### Conclusions

The higher reproductive success of *Ceratina* mothers in sun certainly explains their strong preference for nesting in sun. Despite the costs to juveniles, mothers clearly have higher fitness in sun than in shade in our study populations. However, costs and benefits of nesting in sun and shade could differ from place to place and from time to time. For instance, urban *Ceratina* populations might experience lower predation, which might even up the rates of total nest failure in sun and shade. At the same time, urban heat islands might increase the thermoprotective costs to juveniles of sun nesting (Hamblin et al. 2017). Such patterns might produce a cost/benefit calculus favouring shade-nesting. Climate change also seems very likely to change the fitness calculus; as very hot summer days become more frequent, the cost of thermoprotection in sun nests may rise to the point where sun nesting loses its advantage.

Changes or shifts in thermal breadth are not uncommon in Hymenoptera. Urban honeybees exposed to higher average temperatures shifted thermal breadth (Sánchez-Echeverría et al. 2019), while urban-dwelling acorn-nesting ants that experienced higher temperatures, displayed a small increase in heat tolerance and substantial loss of cold tolerance (Diamond et al. 2018). These patterns raise some interesting questions for future research on *C. calcarata* nesting in sun and shade. How persistent are shifts in thermal breadth of sun bees? Might females born in sun nests be better adapted to nesting in sun themselves? Another question relates to the relationship between CT_max_ and CT_min_. Does a rise in CT_max_ necessitate a rise in CT_min_? If so, then *Ceratina* raised in sun nests might be less cold tolerant and less likely to survive hibernation.

## Acknowledgments

We thank Lyndon Duff for providing R-code for comparing annual temperature patterns, and the entire Brock Bee Lab for numerous discussions. We thank Peter Kryger at Niagara Region Waste Management Division for ensuring our access to field sites even when public sites were closed down. Funding was provided by Brock University Faculty of Graduate Studies to JdH, an NSERC Undergraduate Summer Research Award to JM, and NSERC Discovery Grants to GT and MHR, and is much appreciated.

