## Supplemental tables and figures for "Costs and benefits of maternal nest choice: tradeoffs between brood survival and thermal stress for small carpenter bees"

**Table S1.** BeeCR temperature profiles over 24 hours. In a pilot study in 2020, bees raised in rows 1 and 2 frequently died or failed to reach adulthood, suggesting that these temperatures were too low for pupal development. In 2021, bees were raised in rows 2 to 11 (see Supplemental Figure S3).

| **Hour** | **BeeCR Set point (°C)** | **Temperature of column (°C)** | | | | | | | | | | |
| --- | --- | --- | --- | --- | --- | --- | --- | --- | --- | --- | --- | --- |
| **0:00** | 18.8 | 8.7 | 9.3 | 10.6 | 12.6 | 15.1 | 17.9 | 20.7 | 23.4 | 25.9 | 28 | 29.4 |
| **1:00** | 19 | 8.9 | 9.5 | 10.8 | 12.8 | 15.3 | 18.1 | 20.9 | 23.6 | 26.1 | 28.2 | 29.6 |
| **2:00** | 19 | 8.9 | 9.5 | 10.8 | 12.8 | 15.3 | 18.1 | 20.9 | 23.6 | 26.1 | 28.2 | 29.6 |
| **3:00** | 18.9 | 8.8 | 9.4 | 10.7 | 12.7 | 15.2 | 18 | 20.8 | 23.5 | 26 | 28.1 | 29.5 |
| **4:00** | 19.2 | 9.1 | 9.7 | 11 | 13 | 15.5 | 18.3 | 21.1 | 23.8 | 26.3 | 28.4 | 29.8 |
| **5:00** | 19.1 | 9 | 9.6 | 10.9 | 12.9 | 15.4 | 18.2 | 21 | 23.7 | 26.2 | 28.3 | 29.7 |
| **6:00** | 19 | 8.9 | 9.5 | 10.8 | 12.8 | 15.3 | 18.1 | 20.9 | 23.6 | 26.1 | 28.2 | 29.6 |
| **7:00** | 23.5 | 13.5 | 14 | 15.4 | 17.4 | 19.9 | 22.6 | 25.4 | 28.2 | 30.6 | 32.6 | 34 |
| **8:00** | 23.6 | 13.6 | 14.1 | 15.5 | 17.5 | 20 | 22.7 | 25.5 | 28.3 | 30.7 | 32.7 | 34.1 |
| **9:00** | 23.7 | 13.7 | 14.2 | 15.6 | 17.6 | 20.1 | 22.8 | 25.6 | 28.4 | 30.8 | 32.8 | 34.2 |
| **10:00** | 25 | 15 | 15.5 | 16.9 | 18.9 | 21.4 | 24.1 | 26.9 | 29.7 | 32.1 | 34.1 | 35.5 |
| **11:00** | 25 | 15 | 15.5 | 16.9 | 18.9 | 21.4 | 24.1 | 26.9 | 29.7 | 32.1 | 34.1 | 35.5 |
| **12:00** | 25 | 15 | 15.5 | 16.9 | 18.9 | 21.4 | 24.1 | 26.9 | 29.7 | 32.1 | 34.1 | 35.5 |
| **13:00** | 25.3 | 15.3 | 15.8 | 17.2 | 19.2 | 21.7 | 24.4 | 27.2 | 30 | 32.4 | 34.4 | 35.8 |
| **14:00** | 26.8 | 16.8 | 17.3 | 18.7 | 20.7 | 23.2 | 25.9 | 28.7 | 31.5 | 33.9 | 35.9 | 37.3 |
| **15:00** | 25.8 | 15.8 | 16.3 | 17.7 | 19.7 | 22.2 | 24.9 | 27.7 | 30.5 | 32.9 | 34.9 | 36.3 |
| **16:00** | 26.1 | 16.1 | 16.6 | 18 | 20 | 22.5 | 25.2 | 28 | 30.8 | 33.2 | 35.2 | 36.6 |
| **17:00** | 25.8 | 15.8 | 16.3 | 17.7 | 19.7 | 22.2 | 24.9 | 27.7 | 30.5 | 32.9 | 34.9 | 36.3 |
| **18:00** | 25.2 | 15.2 | 15.7 | 17.1 | 19.1 | 21.6 | 24.3 | 27.1 | 29.9 | 32.3 | 34.3 | 35.7 |
| **19:00** | 21 | 10.9 | 11.5 | 12.9 | 14.9 | 17.3 | 20.1 | 22.9 | 25.6 | 28.1 | 30.2 | 31.5 |
| **20:00** | 20.4 | 10.3 | 10.9 | 12.3 | 14.3 | 16.7 | 19.5 | 22.3 | 25 | 27.5 | 29.6 | 31 |
| **21:00** | 19.6 | 9.5 | 10.1 | 11.4 | 13.5 | 15.9 | 18.7 | 21.5 | 24.2 | 26.7 | 28.8 | 30.2 |
| **22:00** | 19.2 | 9.1 | 9.7 | 11 | 13 | 15.5 | 18.3 | 21.1 | 23.8 | 26.3 | 28.4 | 29.8 |
| **23:00** | 18.9 | 8.8 | 9.4 | 10.7 | 12.7 | 15.2 | 18 | 20.8 | 23.5 | 26 | 28.1 | 29.5 |
| **Average daily temperature** | 22.2 | 12.2 | 12.7 | 14.1 | 16.1 | 18.6 | 21.3 | 24.1 | 26.9 | 29.3 | 31.4 | 32.8 |
|  |  | **1** | **2** | **3** | **4** | **5** | **6** | **7** | **8** | **9** | **10** | **11** |

**Table S2.** Generalized linear model examining variation in high temperature tolerance (CTmax) of brood measured as adults in 2019 and 2020.

| **Predictor Variable** | **NumDF** | **DenDF** | **F value** | **p** |
| --- | --- | --- | --- | --- |
| Dry Weight | 1 | 67.622 | 1.193 | 0.279 |
| Treatment | 1 | 9.529 | 11.863 | **0.007** |
| Year | 1 | 11.424 | 22.881 | **<0.001** |

Notes: Treatment and year effects represent partial effects from generalized linear models after accounting for differences between males and females, with brood cell position and similarities among nestmates treated as random effects. Significant values bolded.

Abbreviations: Sum Sq, sum of squares; Mean Sq, mean squares; NumDF, numerator degrees of freedom; DenDF, denominator degrees of freedom.

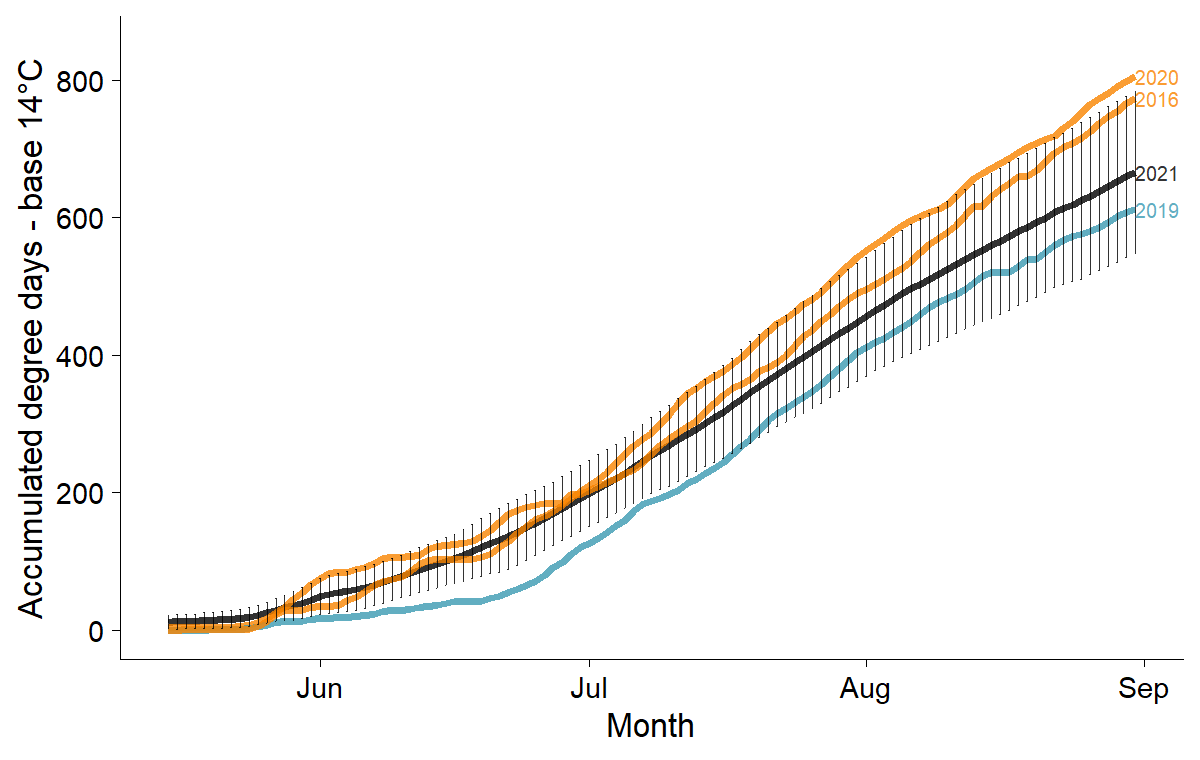

**Figure S1.** Annual heat accumulation during the Ceratina flight seasons of 2016, 2019 and 2021, based on a degree-day threshold of 14°C. The vertical black bars represent the 30 year average as calculated in 2021. Data are from the Environment Canada weather station at Vineland (https://climate.weather.gc.ca/climate_data/daily_data_e.html?StationID=31367). Kolmogorov-Smirnov tests indicate that 2020 was significantly warmer than 2019 (D=0.22, p=0.10).

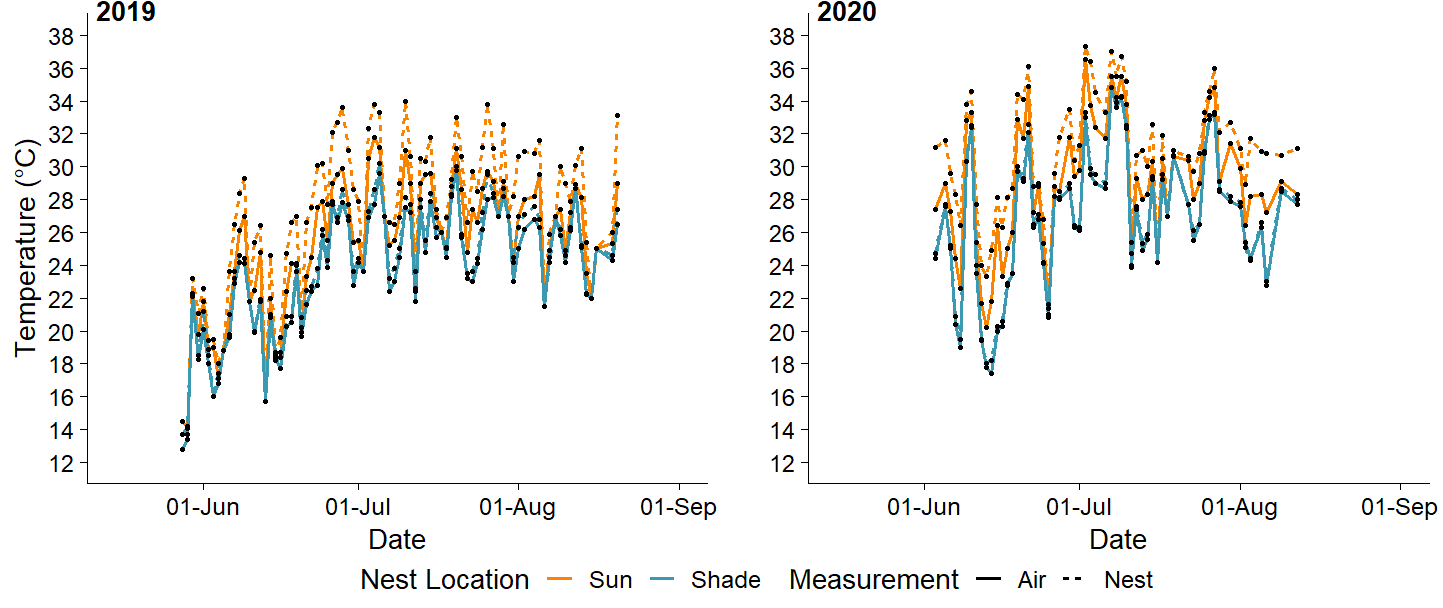

**Figure S2.** Nest and air temperatures measured in sun and shade nesting sites over the brood development period in 2019 and 2020. Nest and air temperatures were measured from 28 May to 20 August 2019 and from 3 June to 12 August 2020. Sun nests were also generally hotter than shade nests when measuring air and nest temperatures at ~2pm.
